# Macrocyclic colibactin induces DNA double-strand breaks via copper-mediated oxidative cleavage

**DOI:** 10.1101/530204

**Authors:** Zhong-Rui Li, Jie Li, Wenlong Cai, Jennifer Y. H. Lai, Shaun M. K. McKinnie, Wei-Peng Zhang, Bradley S. Moore, Wenjun Zhang, Pei-Yuan Qian

## Abstract

Colibactin is an as-yet-uncharacterized human gut bacterial genotoxin, whose biosynthesis is linked to *clb* genomic island that distributes widespread in pathogenic and commensal human enterobacteria. Colibactin-producing gut microbes promote colon tumor formation and enhance progression of colorectal cancer (CRC) via DNA double-strand breaks (DSBs)-induced cellular senescence and death; however, the chemical basis contributing to the pathogenesis at the molecular level remains elusive. Here we report the discovery and the mechanism of action of colibactin-645 as the highly sought final colibactin metabolite with a novel molecular scaffold. Colibactin-645 recapitulates its previously assumed genotoxicity and cytotoxicity, exhibiting a strong DNA DSBs activity *in vitro* and in human cell cultures via a unique copper-mediated oxidative mechanism. We also present a complete model for colibactin biosynthesis, revealing an unprecedented dual function of the aminomalonate-utilizing polyketide synthases. This work thus provides the first molecular basis for colibactin’s genotoxic activity and facilitates further mechanistic study of colibactin-related CRC incidence and prevention.

## Main Text

Human microbiota is a massive consortium of all microbes that reside in and on human bodies. These microbes are increasingly being correlated to human health and disease, but the underlying molecular mechanisms of human-microbe interactions often remain elusive^1,2^. Interrogating the specialized metabolites produced by human microbiota allows a thorough study of chemical regulatory and signaling processes, and improves our understanding of the interplay between microbiota and host at a molecular level. Despite the importance of these small molecules in human health and disease, it is often challenging to characterize them because of the difficulty in the culture and genetics of producing microbes and the low titers of these metabolites^3–5^.

A well-known example of such specialized metabolite is colibactin, a cryptic human gut bacterial genotoxin that has captured the attention of both biologists and chemists due to its significant effects on human health and intriguing biosynthetic logic^6–8^. The biosynthesis of colibactin is linked to a 54-kilobase nonribosomal peptide synthetase (NRPS)-polyketide synthase (PKS) hybrid gene cluster^9^ (*clb* pathogenicity island), which has been phenotypically associated with the pathogenesis of colorectal cancer (CRC). In particular, *in vitro* infection with *Escherichia coli* strains harboring *clb* induced DNA double-strand breaks (DSBs) in cultivated human cells, leading to cell cycle arrest and eventually cell death^9^. Subsequent physiological studies showed that *clb*^+^ bacteria induced *in vivo* DNA damage and genomic instability in enterocytes^10^, caused cellular senescence^11,12^, increased intestinal permeability^13^, and promoted colon tumor formation in mouse models of chronic intestinal inflammation^12,14,15^, suggesting that these bacteria could promote human CRC development on a broader level^8^. Consistently, *clb*^+^ *E. coli* was over-represented in biopsies isolated from CRC patients compared to non-CRC controls (~60% vs. ~20%, respectively)^14,16^. In addition to its remarkable association with human health, the *clb* island was also identified in the genomes of other proteobacteria, including coral and honeybee symbionts, suggesting an even more comprehensive role that colibactin might play in mediating evolutionarily conserved or consistent interactions between bacteria and hosts^17,18^.

Given the physiological importance of intestinal pathology induced by human body’s microscopic residents, it is urgent to reveal the molecular identity of genotoxic colibactin as the missing link between certain gut microbes and DNA DSBs and decode the mechanism underlying colibactin-induced DNA damage. Despite tremendous efforts, colibactin’s structural elucidation remains a formidable challenge due to its instability, low titer, and the elusive and complex biosynthetic logic of *clb* pathway^19–29^. This knowledge gap has prevented comprehensive studies of colibactin-related CRC incidence and prevention, and limited mechanistic investigations of even more extensive influence of *clb* island on microbe-host interactions.

In order to investigate the corresponding genotoxic colibactin that possesses intrinsic DNA DSBs activity and causes chromosome aberrations, the following three issues need to be addressed. 1) The mutation of individual *clb* genes revealed that all genes encoding NRPS-PKS and associated biosynthetic enzymes were indispensable to the genotoxicity of *clb* island^9,26^, however, the final colibactin metabolite that requires all of the *clb* genes for its biogenesis has not been identified. 2) The precise role of ClbP, a membrane-bound peptidase that was proposed to be important for colibactin maturation^19,20^, remains unknown. 3) The induction of DNA DSBs has been defined as a signature feature of *clb* island^6–10^, yet the conclusive evidence for colibactin directly mediating DNA breakage is still lacking, despite that precolibactin-546 (**5**) showed a weak DNA crosslinking activity *in vitro* in the presence of reducing agents^24^ (Fig. 1a). Of the many types of DNA damage that exist within cells, the DNA DSBs are considered to be the most hazardous lesions^30^, suggesting the remarkable cytotoxicity of the yet-to-be-identified colibactin metabolite. Here we report the structural elucidation of the final mature colibactin, and further show that colibactin induces DNA DSBs *in vitro* and in various human cell cultures via a unique copper-mediated oxidative mechanism.

**Fig. 1.**
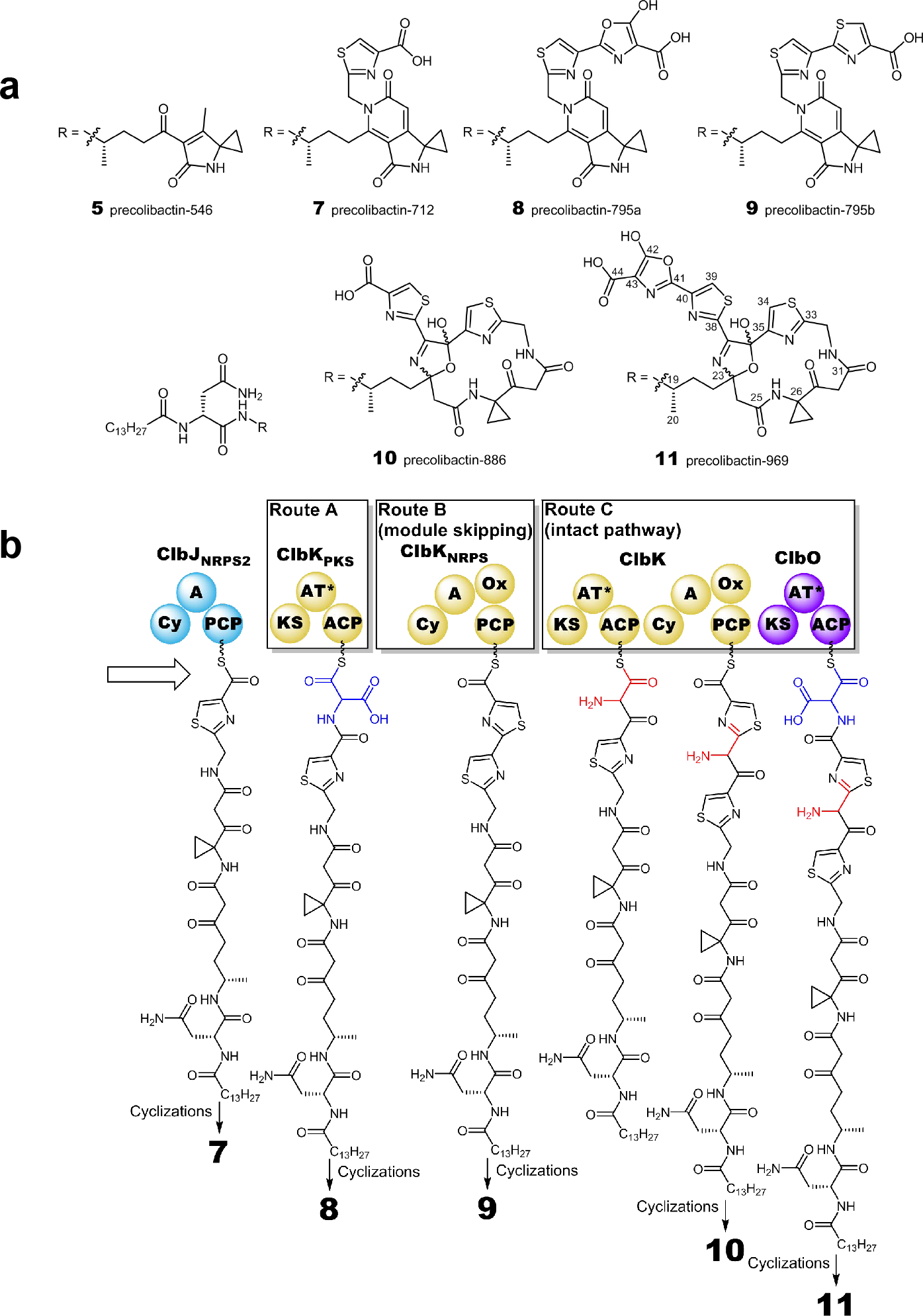
Structures and proposed biosynthesis of precolibactins. **a**, Structures of precolibactin-546 (**5**), precolibactin-712 (**7**), precolibactin-795a (**8**), precolibactin-795b (**9**), precolibactin-886 (**10**) and precolibactin-969 (**11**). **b**, Proposed biosynthetic pathway of precolibactins. Extending from ClbJ, the dimodule PKS/NRPS ClbK shows diverse functions in the production of *clb* metabolites. The *clb* pathway utilizes only ClbK_PKS_ to produce **8** (Route A); or skips ClbK_PKS_ but utilizes ClbK_NRPS_ to produce **9** (Route B); or utilizes both of these two modules to produce **10** which is the precursor for the assembly of **11** (Route C). A, adenylation; ACP, acyl carrier protein; AT, acyltransferase; Cy, cyclization; KS, ketosynthase; Ox, oxidase; PCP, peptidyl carrier protein. AT* domains are predicted based on structural topology as ancestral inactive relics.

## Results

### Discovery of complete colibactin precursor

Our previous efforts to identify colibactin biosynthetic intermediates resulted in the structural elucidation of precolibactin-886 (**10**)^28^, which was isolated from a *clb*^+^ heterologous expression strain *E. coli* DH10B/pCAP01-*clb* with disrupted *clbP* and *clbQ* that encode a peptidase and a type II thioesterase mediating the off-loading of *clb* pathway intermediates, respectively^19,28,31^ (Fig. 1). The double mutation of Δ*clbP*Δ*clbQ* increased the titer of downstream metabolites from the NRPS-PKS assembly line, enabling the structural characterization of **10** whose biogenesis requires all components of the assembly line except the PKS ClbO^28^. We then searched for a more complete colibactin derivative that could account for the activity of ClbO. The initial examination of the Δ*clbP*Δ*clbQ* and Δ*clbP*Δ*clbQ*Δ*clbO* mutants for the selective loss of metabolites identified a precolibactin metabolite with *m/z* 970 (named precolibactin-969, **11**) in a trace amount (Fig. 1, Fig. 2a). To facilitate the structural elucidation of **11**, additional regulatory/resistance *clb* genes including *clbR* and *clbS* were explored to probe their effects on the production of **11**. ClbR is a known positive transcriptional regulator and its overexpression previously led to a five-fold increase in the prodrug motif accumulation^22^, and ClbS is a colibactin resistance protein that was proposed to sequester or modify colibactin and thereby prevent self-inflicted DNA damage^32,33^. While overexpression of *clbR* had no obvious effect on the titer of **11**, inactivation of *clbS* resulted in a notable four-fold increase in the titer of **11** along with other precolibactins (Fig. 2a). The observed eliciting phenomenon in Δ*clbS* is consistent with the proposed function of ClbS, and we thus used the Δ*clbP*Δ*clbQ*Δ*clbS* mutant strain for the subsequent precolibactin production and purification.

**Fig. 2.**
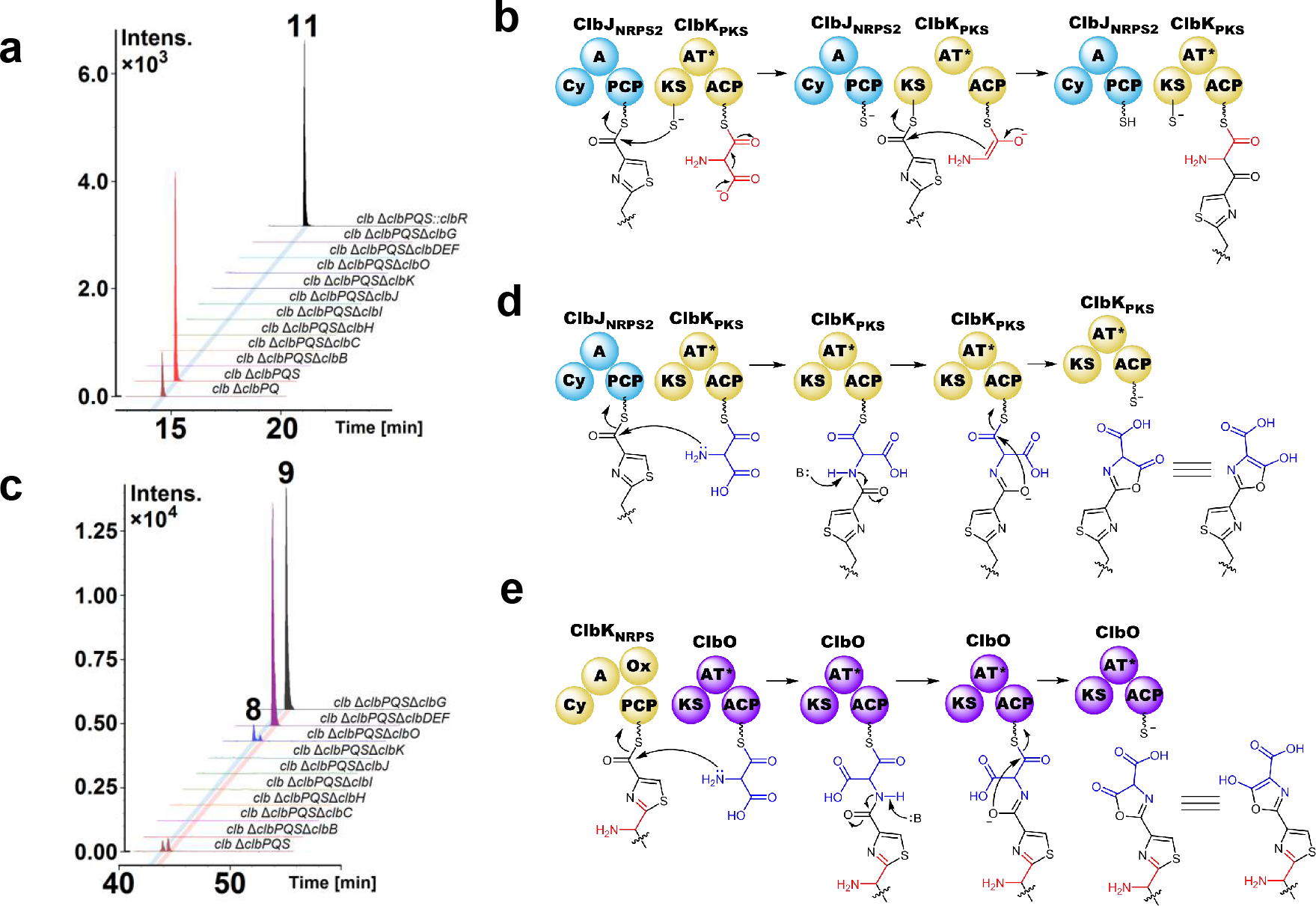
Genes and proposed mechanisms of aminomalonate-utilizing PKSs in the biosynthesis of precolibactins. **a**, A comparison of LC–MS extracted ion chromatogram traces of the metabolic extracts from Δ*clbP*Δ*clbQ*, Δ*clbP*Δ*clbQ*Δ*clbS* and its nine mutants, and Δ*clbP*Δ*clbQ*Δ*clbS::clbR*, showing the impact of gene knockout or knockin on the yield of **11**, and the requirement of *clb* pathway genes for the biosynthesis of **11**. EIC+ = 970.38 ± 0.01, which corresponds to **11**. **b**, Proposed mechanism of ClbK_PKS_ underlying the production of **10.** The chain elongation is achieved through C−C bond formation by decarboxylative Claisen condensation. **c**, A comparison of LC–MS extracted ion chromatogram traces of the metabolic extracts from *clbP*Δ*clbQ*Δ*clbS* and its nine mutants. EIC+ = 796.37 ± 0.01 and 796.35 ± 0.01, which correspond to **8** and **9**, respectively. **d**, Proposed mechanism of ClbK_PKS_ underlying the production of **8**. The chain elongation is achieved through C−N bond formation by nucleophilic attack of the amine in the aminomalonate extender unit, followed by synchronous cyclization and release of **8**. **e**, Proposed mechanism of ClbO underlying the production of **11** with a similar biosynthetic logic to that of **8**. **b**, **d**and **e**, A, adenylation; ACP, acyl carrier protein; AT, acyltransferase; Cy, cyclization; KS, ketosynthase; Ox, oxidase; PCP, peptidyl carrier protein. AT* domains are predicted based on structural topology as ancestral inactive relics.

From a 2,000-L fermentation culture of Δ*clbP*Δ*clbQ*Δ*clbS*, 50 μg of **11** was obtained after extraction with organic solvent followed by multiple rounds of reversed-phase liquid chromatography purification. **11** was isolated as white and amorphous powder, and its molecular formula was determined as C_44_H_59_N_9_O_12_S_2_ by high-resolution mass spectrometry (HRMS) (*m*/*z* 970.3799, calculated: 970.3797), which has an additional C_3_HNO_2_ compared to the formula of **10**. The presence of an extra nitrogen atom in **11** is consistent with the known aminomalonate substrate utilization by ClbO^26,27^, which was also supported by the isotope-labeled precursor feeding experiments, suggesting the incorporation of an additional aminomalonate compared to **10**. Similar to **10**, **11** was isolated as an approximately equal mixture of two isomers. Analysis of extensive nuclear magnetic resonance (NMR) spectra and high-resolution tandem mass spectrometry (HRMS^n^) fragmentation data demonstrated that **11** and **10** share the same macrocyclic scaffold from C-1 to C-40 (Fig. 1), indicating that ClbO functions towards the end of the NRPS-PKS assembly line to incorporate the last building monomer of aminomalonate. However, we were not able to assign the structure of this extra region (C-41 to C-44) based on the NMR spectra due to the apparent proton deficiency feature and the extremely low titer of **11** at this stage.

We then turned to the PKS activity of ClbO to predict the fate of the corresponding aminomalonate unit. In the *clb* locus, two PKS modules, ClbK_PKS_ and ClbO, were enzymatically established to incorporate an aminomalonate extender unit^26,27^. Both PKS modules have domains organized into KS-AT*-ACP (Fig. 1b). A maximum likelihood tree revealed a close phylogenetic relationship between these two KS domains, suggesting a similar activity of ClbK_PKS_ and ClbO. While ClbK_PKS_ was shown to incorporate aminomalonate through a decarboxylative Claisen condensation in forming **10** (Fig. 2b), this reactivity does not account for the addition of three carbon atoms promoted by ClbO in forming **11**. Considering the typical observation that the titers of upstream colibactin metabolites were significantly higher than those of downstream metabolites^25,28^, we searched for a possible intermediate that is stalled at ClbK_PKS_ with an additional of C_3_HNO_2_ in its molecular formula compared to precolibactin-712 (**7**) to facilitate the total structural determination of **11** (Fig. 1). Careful analysis of the culture extracts of Δ*clbP*Δ*clbQ*Δ*clbS*Δ*clbO* revealed a new metabolite (named precolibactin-795a, **8**) with the molecular formula of C_39_H_53_N_7_O_9_S_1_ (*m*/*z* 796.3697, calculated 796.3698) (Fig. 2c). A total of 1.1 mg of **8** from a 500-L fermentation culture were obtained and extensive analysis of the NMR spectra and HRMS^n^ fragmentation data indicated that in comparison with **7** and precolibactin-795b (**9**), **8** contains a unique 5-hydroxy oxazole moiety next to the terminal carboxyl group (Fig. 1). We propose that to assemble **8**, the aminomalonate unit is incorporated into the assembly line by ClbK_PKS_ through nucleophilic attack of the amine in the aminomalonate extender unit on the upstream peptidyl-*S*-T thioester of ClbJ, followed by synchronous cyclization and release (Fig. 1, Fig. 2d). This novel biosynthetic logic of accommodating a rare aminomalonate building block by PKS was further supported by the gene inactivation and isotope labeled precursor feeding experiments (Fig. 2c). We thus deduce that **11** contains the same 5-hydroxy oxazole moiety next to its terminal carboxyl group, which is derived from the aminomalonate extender unit of ClbO and formed through the same chemical logic as in **8** (Fig. 1, Fig. 2e). The discovery of **8** suggests the dual function of aminomalonate-utilizing PKSs in promoting both the C−C and C−N bond formations in colibactin biosynthesis. Indeed, a precolibactin metabolite (precolibactin-943, **12**) with *m/z* 944 corresponding to the decarboxylative condensation activity of ClbO was also observed, but its titer was only approximately 10% of that of **11**.

### Maturation of colibactin

Precolibactin-969 (**11**) is hitherto the largest colibactin derivative that requires all components of the NRPS-PKS assembly line for its biosynthesis. We next examined whether ClbP, the dedicated peptidase for colibactin maturation, is capable of hydrolyzing this precursor in the bacterial periplasm and releasing the mature colibactin (Fig. 3a). Incubation of **11** with the culture of *E. coli* expressing ClbP resulted in the complete loss of **11** and the production of both the prodrug motif *N*-myristoyl-D-asparagine (**14**) and a new metabolite (named colibactin-645, **13**) with the molecular formula of C_26_H_27_N_7_O_9_S_2_ (*m*/*z* 646.1394, calculated 646.1384) (Fig. 3). **13** was confirmed to be the mature compound of **11** with a free *N*-terminus after cleavage and release of the prodrug motif based on the comparative HR-MS/MS analysis. It is notable that different from **11** and **14**, **13** is a very water-soluble compound which could not be extracted by typical organic solvents such as ethyl acetate^21,24^. Additionally, we observed a significantly increased recovery yield of **13** from the ClbP-expressing *E. coli* culture upon treatment of metal chelators, such as ethylenediaminetetraacetic acid (EDTA) and Chelex-100 (Fig. 3b). The positive effect of metal chelators on metabolite yields from *E. coli* cultures was also observed for **11**, but not for other precolibactins such as **2**, **5**, and **7**. These results suggested the susceptibility of colibactin-645 (**13**) and its precursor (**11**) to trace metals for possible degradation.

**Fig. 3.**
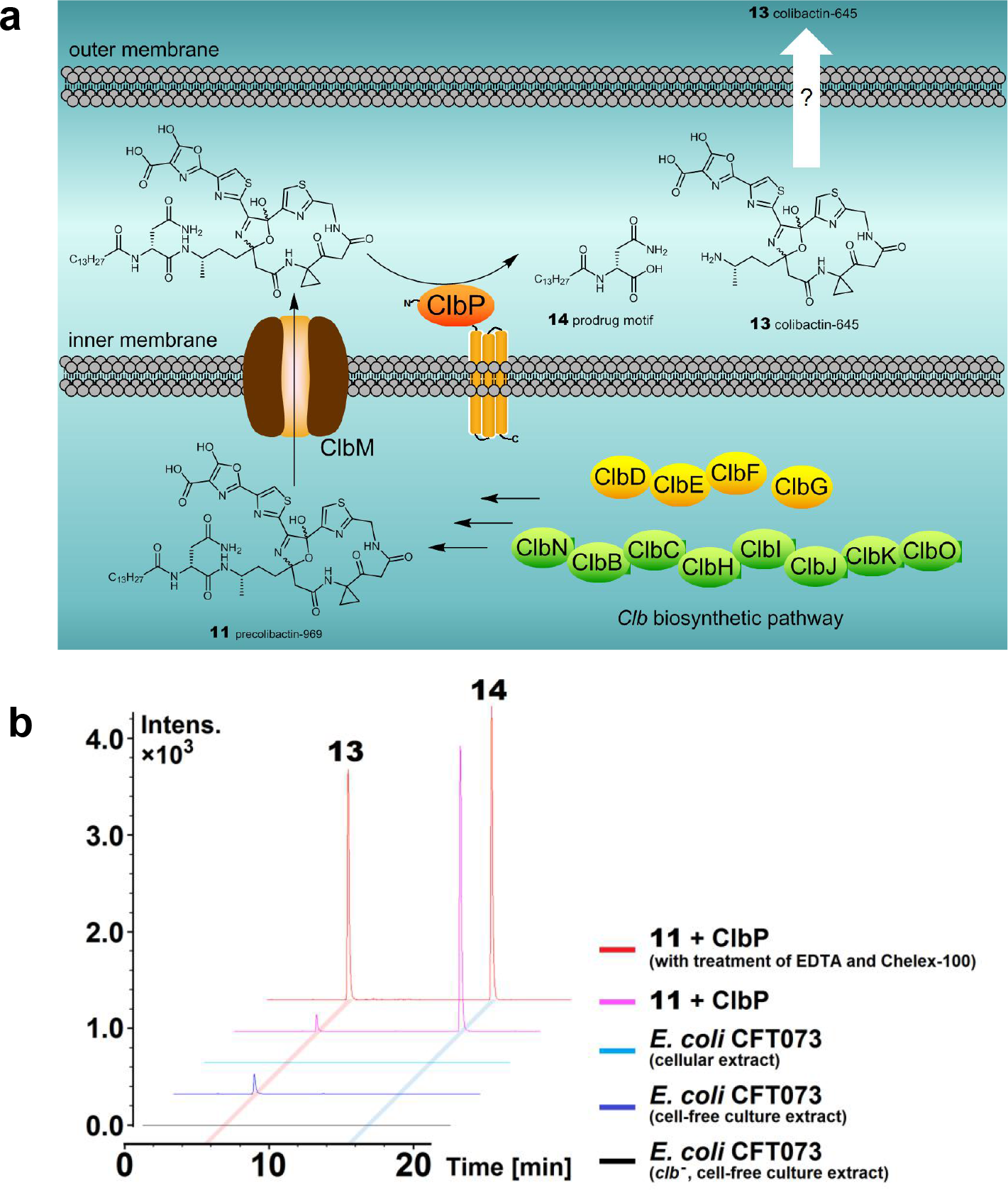
Maturation of colibactin. **a**, Proposed pathway for colibactin maturation. A prodrug mechanism is involved in colibactin biosynthesis. Precolibactin-969 (**11**) is biosynthesized in the cytoplasm of *E. coli* strains by the *clb* biosynthetic pathway and transported via ClbM into the periplasm, whereby the membrane-bound peptidase, ClbP, cleaves **11** to generate mature colibactin-645 (**13**) and a prodrug motif *N*-myristoyl-D-asparagine (**14**), followed by outer membrane translocation. **b**, A comparison of LC-MS extracted ion chromatogram traces shows the production of **13** resulting from its precursor **11** cleavage by *E. coli* strains expressing the peptidase gene *clbP* in the presence or absence of metal chelators; and the detection of metabolite identical to **13** from either cell-free culture extracts or cellular extracts of cultured wild-type *clb*^+^ *E. coli* CFT073 and its *clb*^−^ mutant. EIC+ = 646.14 ± 0.01 and 343.26 ± 0.01, which correspond to **13** and **14**, respectively.

### Colibactin production by a native strain

We next investigated whether the native *clb*^+^ *E. coli* strain could produce the same colibactin-645 to probe if **13** was a native metabolite or an artifact arising from a non-natural biosynthetic pathway in a heterologous host. LC-MS analysis of cell-free culture extracts of the wild-type *clb*^+^ *E. coli* CFT073 and its *clb*^−^ mutant revealed a peak identical to **13** only in the wild-type *clb*^+^ strain, confirming that **13** is the native product of the *clb* pathogenicity island (Fig. 3b). It is notable that after enrichment from a 2-L of fermentation culture, only a trace amount of **13** was detected by HRMS analysis, indicating the low titer of **13** or its chemical lability. Since previous work showed that direct contact between bacterial and eukaryotic cells was required for full toxicity of colibactin^9^, we further examined whether a majority of **13** are associated with the producing cells. **13** was not detected in the cellular extract of *clb*^+^ *E. coli* CFT073 (Fig. 3b), suggesting that the mature colibactin was secreted after production and highly unstable after secretion.

### DSBs activity of colibactin *in vitro*

After obtaining the highly sought mature colibactin (**13**), we examined its DNA DSBs *in vitro* using the pBR322 plasmid DNA strand scission assay, a surrogate test for DNA damage^34^. Although **13** showed sparse DNA damage activity upon incubation with DNA, in the presence of Cu(II), but not other metals such as Fe(III) and Fe(II), both **13** and its precursor precolibactin-969 (**11**) caused significant DNA breakage with the formation of both nicked (Form II) and linearized (Form III) DNA from the supercoiled plasmid DNA (Form I) (Fig. 4a). Since **13** and **11** demonstrated comparable DNA damage activities in initial tests, we used **11** as an appropriate substitute for **13** in the following *in vitro* assays because **11** was more readily available. A time-course experiment of DNA cleavage was then performed to determine if the colibactin-induced linearized DNA arose from coupled strand-cleavage events (DSBs), or from an accumulation of unrelated single-strand breaks (SSBs). All three forms of DNA were visible on the gel, showing classical evidence of DSBs (Fig. 4b). A Freifelder–Trumbo analysis was further performed to calculate the number of SSBs (*n*_1_) and DSBs (*n*_2_) per molecule of DNA after treatment with **11** at various time points, which resulted in a constant ratio of SSBs to DSBs (5.35:1). This number is significantly lower than 120:1 that was expected if DSBs were to arise from an accumulation of unrelated SSBs^35^, and is comparable to some of the well-known DNA DSBs inducers including (–)-lomaiviticin A (5.3:1) and bleomycin (9:1)^34,36^, supporting the coupled strand-cleavage activity of colibactin. It is notable that under the same reaction condition, **11** displayed a stronger DNA DSBs activity than bleomycin which also requires the presence of a redox-active metal ion for DNA cleavage^37,38^.

**Fig. 4.**
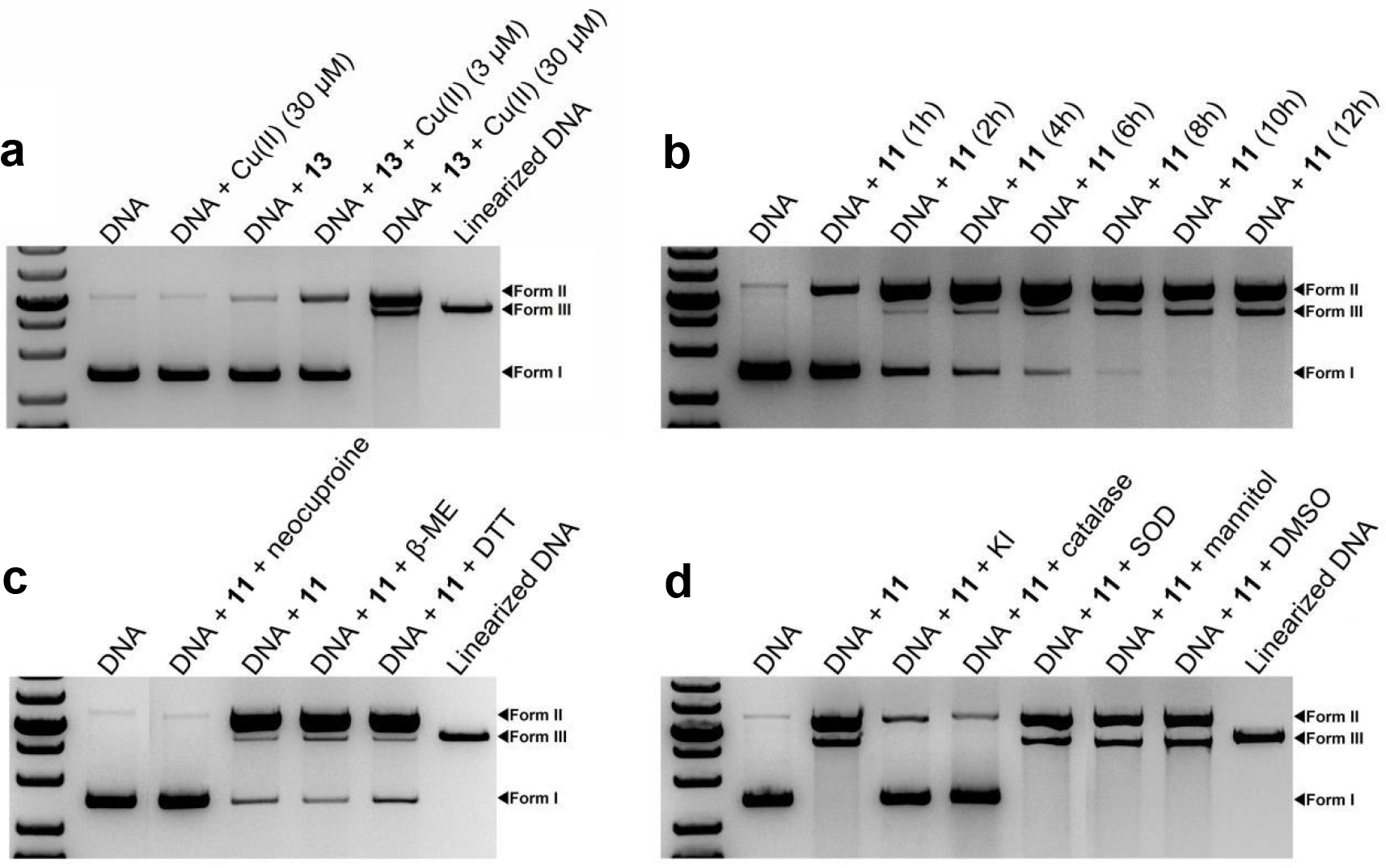
Analysis of DNA damage by colibactin *in vitro*. **a**, The effect of colibactin-645 (**13**) on the plasmid pBR322 DNA cleavage. Reactions were performed at 15 μM **13** in the absence or presence of Cu(II) (3 μM or 30 μM) for 12 hours at 37 °C. DNA cleavage by **13** is observed only in the co-incubation of Cu(II) and **13**, in which nicked (Form II) and linearized (Form III) DNA forms from the supercoiled plasmid DNA (Form I). **b**, The time-dependent DNA damage induced by precolibactin-969 (**11**) (15 μM) is observed in the presence of Cu(II) (30 μM). Reactions were performed at 37 °C with different incubation times. **c**, The effect of a specific Cu(I) chelator neocuproine (1 mM), a reductant β-mercaptoethanol (β-ME) (5 mM), or a reductant dithiothreitol (DTT) (5 mM) on the DNA cleavage by **11** (15 μM) in the presence of Cu(II) (30 μM). Reactions were performed at 37 °C for 4 h. **d**, The effect of various reactive oxygen species (ROS) scavengers, including potassium iodide (KI) (1 mM), catalase (0.1 mg/mL), superoxide dismutase (SOD) (10 units), mannitol (50 mM), and dimethyl sulfoxide (DMSO) (10%), on the **11**-induced DNA cleavage in the presence of Cu(II) (30 μM). Reactions were performed at 15 μM **11**, 37 °C for 12 h. **b**, **c**and **d**, All of the controls (reactions without **11**) of each reagent or scavenger show no DNA cleavage similar to the negative control presented in the figure (the lane with DNA only). **a**–**d**, Top band, nicked DNA (Form II); middle band, linearized DNA (Form III); bottom band, supercoiled DNA (Form I). *Eco*RI-linearized pBR322 DNA is shown as the linearized DNA standard.

The observed Cu(II)-mediated DSBs activity of colibactin is reminiscent of the oxidative mechanism of DNA cleavage involving a metal center reduction^35,39^. The addition of neocuproine, a specific Cu(I) chelator, completely sequestered the DSBs activity of **11**, suggesting that Cu(I) is an essential component for **11**-induced DNA cleavage (Fig. 4c). Surprisingly, the presence of a reducing agent, such as β-mercaptoethanol (β-ME) or dithiothreitol (DTT), had no obvious effect on the DSBs activity of **11** (Fig. 4c). We thus propose that the reduction of Cu(II) to Cu(I) may be mediated by the DNA or by **11** itself, and the latter was supported by the free Cu(I) determination assays upon incubation of **11** and Cu(II). In addition, **10** demonstrated a comparable copper reduction activity as **11**, suggesting that the same macrocyclic scaffold in both compounds could be the active center for Cu(II) binding and reduction. The parallel monitor of the mixture of **10** and Cu(II) by HRMS further showed a loss of the mass signal for **10** over time which was accompanied by an approximately stoichiometric formation of Cu(I), and also a presence of a new mass signal with an isotopic pattern of copper-bound complex^40^. Although this new mass signal was weak and transient which prevented its further characterization, this data supported the direct binding of **10** to copper and the instability of **10** in the presence of copper.

The oxidative mechanism of DNA cleavage was further probed by adding various reactive oxygen species (ROS) scavengers. Plasmid DNA damage by **11** was not measurably influenced by the hydroxyl radical scavengers mannitol and dimethyl sulfoxide (DMSO) (Fig. 4d), which argues against participation of the freely diffusible hydroxyl radical in the observed cleavage and distinguishes the mechanism by which colibactin incises DNA from a sole Fenton-like one^41^. The addition of superoxide dismutase (SOD), which catalyzes the conversion of the superoxide radical into hydrogen peroxide (H_2_O_2_), did not measurably influence DNA cleavage by **11** (Fig. 4d). In contrast, potassium iodide (KI), a H_2_O_2_ scavenger, and catalase, which mediates the decomposition of H_2_O_2_, significantly inhibited the cleavage reaction (Fig. 4d). These results suggested that H_2_O_2_ was involved in mediating DNA cleavage *in vitro*, consistent with the observation of a significant increase in H2-DCFDA fluorescence (a sensor of hydroxyl and peroxyl radicals, and hydrogen peroxide production) in non-transformed human lung fibroblast cells infected by colibactin-producing *E. coli*^11^.

The DNA DSBs activity of **11** was next compared to other precolibactins for a preliminary structure–activity relationship study. Under the same reaction condition, **10** displayed a significantly weaker DSBs activity than **11**, demonstrating that the extra 5-hydroxy oxazole moiety in **11** was important for augmenting the DSBs activity. The DSBs activity of **5**, a precolibactin that has previously demonstrated DNA-crosslinking activity due to its aza-spirocyclopropane warhead, was also tested^24^. **5** did not display DNA-damaging activity even at concentrations as high as 5 mM.

### DSBs activity of colibactin in cells

We next examined the DNA damaging activity of colibactin in various human cell lines. Production of phosphorylated histone H2AX (γH2AX) and translocation of the p53 binding protein 1 (53BP1) are early events in the cellular response to DNA DSBs^42,43^. Four hours after exposure to 50 nM of **13**, HeLa cells showed formation and colocalization of foci derived from γH2AX and 53BP1 (Fig. 5a). By comparison, the γH2AX and 53BP1 foci were undetectable in cells treated with 50 nM of **11** (Fig. 5a), in contrast to the comparable activity of **13** and **11** in the pBR322 plasmid DNA strand scission assay. This result supported that maturation was a prerequisite for colibactin’s genotoxicity *in vivo*^15^. In addition, **15**, the mature product of **10** after ClbP cleavage, also demonstrated a significantly lower activity than that of **13** (Fig. 5a), consistent with the lower DSBs activity of **10** than **11** *in vitro*. The similar foci formation and colocalization were also observed in other cell lines such as human normal colon epithelial FHC cells, human normal colon fibroblast CCD-112 CoN cells, and colorectal cancer HCT-116 cells treated with 50 nM of **13**, which established that the cellular response to **13** was not cell-line specific, consistent with previously reported cytopathic effect in various cell lines that were infected by *clb*^+^ *E. coli* strains^9^.

**Fig. 5.**
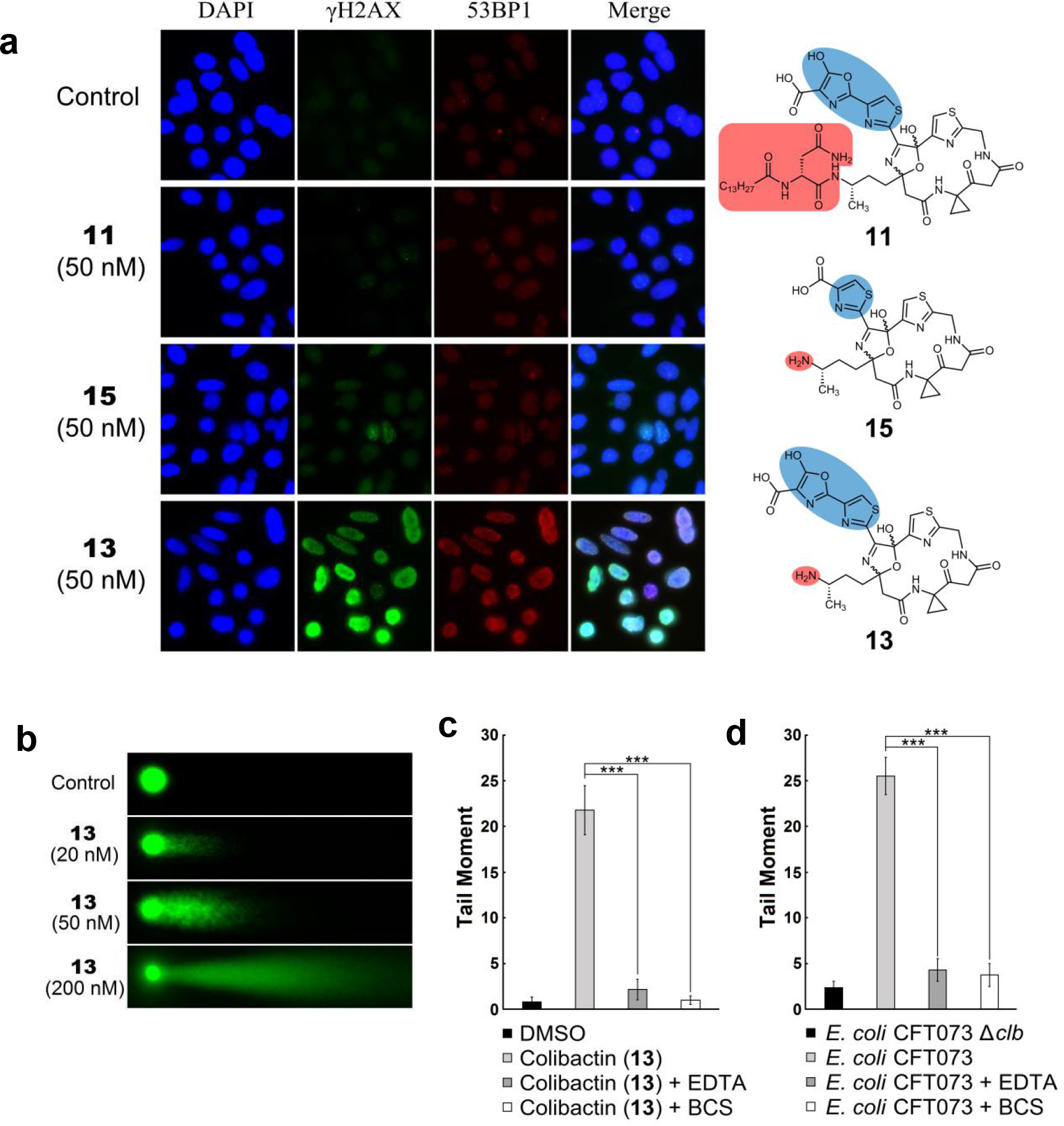
Colibactin-induced DNA damage in cell cultures. **a**, Immunofluorescence imaging of γH2AX and 53BP1 foci in HeLa cells that are treated with precolibactin-969 (**11**, 50 nM), colibactin-645 (**13**, 50 nM) or **15** (50 nM). Columns from left to right, nucleus (blue), γH2AX (green), 53BP1 (red), and merge. In control, only DMSO solvent was added. **b**, Accrued DNA lesions are induced by increased concentrations of **13**, as measured by the neutral comet unwinding assay. **c**, The effect of either ethylenediaminetetraacetic acid (EDTA) (2.5 mM) or bathocuproinedisulfonic acid (BCS) (2 mM) on the DNA damage in HeLa cells after incubation with **13** (50 nM), as measured by the neutral comet assay. **d**, The effect of EDTA (2.5 mM) or BCS (2 mM) on the DNA damage in HeLa cells after incubation with the wild-type *clb*^+^ *E. coli* CFT073, as measured by the neutral comet assay. **c**, **d**, Tail moment was obtained in the neutral comet unwinding assay, which represents the extent of DNA cleavage and is defined as the product of the tail length and the fraction of DNA in the tail. Bars represent mean tail moment (50 cells were randomly selected), error bars represent s.e.m‥ \*\*\**P* < 0.001 (one-way ANOVA).

A neutral comet unwinding assay was also conducted as an effective and independent method to evaluate the occurrence of DNA DSBs in cells treated with **13**^44^. Consistent with the results of γH2AX and 53BP1 induction, a four-hour exposure of Hela cells to **13** caused accrued DNA lesions in a concentration-dependent manner, demonstrated by the migration of cleaved DNA fragments (comet tail) from the nucleoid (comet head) under the influence of an electric field (Fig. 5b). Furthermore, the treatment of either EDTA or bathocuproinedisulfonic acid (BCS), an extracellular Cu-sequestering agent, significantly alleviated the levels of DNA damage caused by the purified compound **13** or the infection of *clb*^+^ *E. coli* CFT073 (Fig. 5c, d), which is in agreement with the observed dependence of copper for colibactin-induced DNA DSBs *in vitro*.

## Discussion

Despite extensive studies on the biology of the *clb* pathogenicity island and the chemistry of the *clb* encoding enzymes, the genotoxic colibactin metabolite with intrinsic DNA DSBs activity had escaped all screening surveillance in the past decade. For the first time, through strain engineering, large-scale fermentation and metabolite comparison, we have identified and characterized the highly sought genotoxic colibactin metabolite, colibactin-645 (**13**). The biosynthesis of **13** requires all predicted biosynthetic enzymes encoded on the *clb* pathogenicity island; more importantly, **13** recapitulates its pre-assumed DNA DSBs activity both *in vitro* and in cell cultures, distinguishing **13** from all previously identified metabolites associated with this pathogenicity island. Considering that all predicted biosynthetic genes were indispensable to the genotoxicity of *clb* island^9,26^, **13** is predicted to be the final mature colibactin metabolite of biological relevance. Interestingly, although macrocyclic colibactins, including **10**, **11**, **13**, and **15**, required copper for their bioactivity, they quickly degraded in the presence of copper, which prevented direct characterization of any colibactin**·**Cu complex. This is akin to the instability of the activated bleomycin that was suggested to have a half-life of only several minutes at 4 °C after binding to a reduced transition metal^45^. In addition to the low abundance and chemical lability, the macrocyclic mature colibactin appeared to be polar compound that stayed in the aqueous phase during organic solvent extraction, which could further contribute to the difficulty in the genotoxic metabolite detection.

The biosynthesis of **13** features a new fate for the atypical aminomalonyl extender unit utilized by PKSs. The incorporation of this aminomalonyl extender unit has been previously elucidated through a traditional decarboxylative Claisen condensation in zwittermicin, guadinomine and colibactin biosynthesis^28,46,47^. In particular, ClbK_PKS_ has been shown to promote the decarboxylative condensation of the aminomalonyl unit that contributes for thiazole and 2,5-dihydro-5-hydroxyoxazole formation in **10** biosynthesis^28^. By comparison, ClbO, the tri-domain PKS that is highly homologous to ClbK_PKS_, has been demonstrated here to preferentially catalyze the non-decarboxylative condensation via amide bond formation that enables the terminal 5-hydroxy oxazole acid generation in **11** and **13** biosynthesis. Furthermore, identification of minor precolibactin metabolites, such as **8** and **12**, indicates that both ClbK_PKS_ and ClbO are capable of facilitating both the C−C and C−N bond formation, demonstrating for the first time the dual function of aminomalonate-utilizing PKSs. It is yet to be determined the molecular basis for these PKSs to prefer one mechanism over the other in producing **13** as the major genotoxic metabolite.

Based on the DNA damage assays both *in vitro* and in cells, we propose the following mechanism for copper-mediated DNA DSBs by colibactin-645 (**13**). After being secreted from a producing bacterium that localizes close to or in contact with the intestinal brush border^10^, **13** binds to exchangeable copper in the intestinal lumen, likely coming from diet^48^, to form a colibactin**·**Cu(II) complex. This complex is quickly transported into the epithelial cell while reduced to a colibactin**·**Cu(I) complex, and the coordination of O_2_ to this cuprous complex in cells generates ‘activated colibactin’ that attacks DNA and initiates DNA cleavage. Cu(II)─O^•^ (or Cu(III)═O) is proposed to be the active species in the ‘activated colibactin’ complex susceptible of DNA carbon–hydrogen bond activation^39^, which is consistent with the observed inhibitory effects of H_2_O_2_ scavengers on the DNA cleavage reaction *in vitro* as colibactin**·**Cu(II)─OOH is a key intermediate to colibactin**·**Cu(II)─O^•^. Additionally, we do not exclude the possibility that **13** quickly enters the epithelial cell and then binds the intracellular copper to exert its activity. This mechanism is analogous to the proposed one for the generation of ‘activated bleomycin’ *in vivo*, differing mainly in the metal usage and the intrinsic metal reduction activity of compounds^37,38^.

The unusual heterocycle-fused macrocycle in **13** is important for copper binding and reduction, as only macrocyclic colibactins, such as **10** and **11**, demonstrated a strong and comparable Cu(II) reduction activity. In addition, the comparison between the DSBs activity of **10** and **11**, as well as **15** and **13**, highlights the significance of the terminal 5-hydroxy oxazole moiety for DNA DSBs activity. We speculate that the thiazole/5-hydroxy oxazole tail found in **11** and **13** may serve as the DNA intercalating element, similar to the function of the bithiazole moiety found in bleomycin^37,38^. Based on the comparative DSBs activity of **11** and **13** *in vitro* but a drastically different solubility as well as a significantly lower activity of **11** in cellular assays, we further propose that the loss of the *N*-terminal fatty acyl-asparagine residue as the prodrug motif facilitates the access of mature colibactin-645 to target eukaryotic cells^15^. Although many secondary metabolites have been reported to induce DNA DSBs, a majority of them function via indirect mechanisms (such as by inhibiting topoisomerase complexes^49^), and few of them cleave DNA double-strand directly^50^. **13** thus represents a novel molecular scaffold exerting a direct DNA DSBs activity, providing a model for designing and synthesizing potent DNA cleaving agents, from synthetic restriction ‘enzymes’ to chemotherapeutic agents.

In summary, we have identified and characterized the highly sought mature genotoxic colibactin metabolite, provided the conclusive evidence for macrocyclic colibactin directly mediating DNA damage, and shed light on the long-standing mystery of the molecular mechanism underlying colibactin-induced DNA DSBs. Our discoveries thus lay out a framework for future investigations that could enhance our understanding of the *clb* pathogenicity island from human gut microbes, and enable further mechanistic interrogation of colibactin-induced DNA DSBs and colibactin-related CRC incidence and prevention.

## References

Nicholson, J. K. et al. Host-gut microbiota metabolic interactions. Science 336, 1262–1267 (2012).

Cho, I. & Blaser, M. J. The human microbiome: at the interface of health and disease. Nat. Rev. Genet. 13, 260–270 (2012).

Sharon, G. et al. Specialized metabolites from the microbiome in health and disease. Cell Metab. 20, 719–730 (2014).

Donia, M. S. et al. A systematic analysis of biosynthetic gene clusters in the human microbiome reveals a common family of antibiotics. Cell 158, 1402–1414 (2014).

Donia, M. S. & Fischbach, M. A. Small molecules from the human microbiota. Science 349, 1254766 (2015).

Bode, H. B. The microbes inside us and the race for colibactin. Angew. Chem. Int. Ed. Engl. 54, 10408–10411 (2015).

Balskus, E. P. Colibactin: understanding an elusive gut bacterial genotoxin. Nat. Prod. Rep. 32, 1534–1540 (2015).

Faïs, T., Delmas, J., Barnich, N., Bonnet, R. & Dalmasso, G. Colibactin: more than a new bacterial toxin. Toxins 10, e151 (2018).

Nougayrède, J. P. et al. Escherichia coli induces DNA double-strand breaks in eukaryotic cells. Science 313, 848–851 (2006).

Cuevas-Ramos, G. et al. Escherichia coli induces DNA damage in vivo and triggers genomic instability in mammalian cells. Proc. Natl. Acad. Sci. U.S.A. 107, 11537–11542 (2010).

Secher, T., Samba-Louaka, A., Oswald, E. & Nougayrède, J. P. Escherichia coli producing colibactin triggers premature and transmissible senescence in mammalian cells. PLOS ONE 8, e77157 (2013).

Cougnoux, A. et al. Bacterial genotoxin colibactin promotes colon tumour growth by inducing a senescence-associated secretory phenotype. Gut 63, 1932–1942 (2014).

Payros, D. et al. Maternally acquired genotoxic Escherichia coli alters offspring’s intestinal homeostasis. Gut Microb. 5, 313–325 (2014).

Arthur, J. C. et al. Intestinal inflammation targets cancer-inducing activity of the microbiota. Science 338, 120–123 (2012).

Tomkovich, S. et al. Locoregional effects of microbiota in a preclinical model of colon carcinogenesis. Cancer Res. 77, 2620–2632 (2017).

Buc, E. et al. High prevalence of mucosa-associated E. coli producing cyclomodulin and genotoxin in colon cancer. PLOS ONE 8, e56964 (2013).

Bondarev, V. et al. The genus Pseudovibrio contains metabolically versatile bacteria adapted for symbiosis. Environ. Microbiol. 15, 2095–2113 (2013).

Engel, P., Vizcaino, M. I. & Crawford, J. M. Gut symbionts from distinct hosts exhibit genotoxic activity via divergent colibactin biosynthesis pathways. Appl. Environ. Microbiol. 81, 1502–1512 (2015).

Brotherton, C. A. & Balskus, E. P. A prodrug resistance mechanism is involved in colibactin biosynthesis and cytotoxicity. J. Am. Chem. Soc. 135, 3359–3362 (2013).

Bian, X. In vivo evidence for a prodrug activation mechanism during colibactin maturation. ChemBioChem 14, 1194–1197 (2013).

Vizcaino, M. I., Engel, P., Trautman, E. & Crawford, J. M. Comparative metabolomics and structural characterizations illuminate colibactin pathway-dependent small molecules. J. Am. Chem. Soc. 136, 9244–9247 (2014).

Brotherton, C. A., Wilson, M., Byrd, G. & Balskus, E. P. Isolation of a metabolite from the pks island provides insights into colibactin biosynthesis and activity. Org. Lett. 17, 1545–1548 (2015).

Bian, X., Plaza, A., Zhang, Y. & Müller, R. Two more pieces of the colibactin genotoxin puzzle from Escherichia coli show incorporation of an unusual 1-aminocyclopropanecarboxylic acid moiety. Chem. Sci. 6, 3154–3160 (2015).

Vizcaino, M. I. & Crawford, J. M. The colibactin warhead crosslinks DNA. Nat. Chem. 7, 411–417 (2015).

Li, Z.-R. et al. Critical intermediates reveal new biosynthetic events in the enigmatic colibactin pathway. ChemBioChem 16, 1715–1719 (2015).

Brachmann, A. O. et al. Colibactin biosynthesis and biological activity depends on the rare aminomalonyl polyketide precursor. Chem. Commun. 51, 13138–13141 (2015).

Zha, L., Wilson, M. R., Brotherton, C. A. & Balskus, E. P. Characterization of polyketide synthase machinery from the pks island facilitates isolation of a candidate precolibactin. ACS Chem. Biol. 11, 1287–1295 (2016).

Li, Z.-R. et al. Divergent biosynthesis yields a cytotoxic aminomalonate-containing precolibactin. Nat. Chem. Biol. 12, 773–775 (2016).

Zha, L. et al. Colibactin assembly line enzymes use S-adenosylmethionine to build a cyclopropane ring. Nat. Chem. Biol. 13, 1063–1065 (2017).

Khanna, K. K. & Jackson S. P. DNA double-strand breaks: signaling, repair and the cancer connection. Nat. Genet. 27, 247–254 (2001).

Guntaka, N. S., Healy, A. R., Crawford, J. M., Herzon, S. B. & Bruner, S. D. Structure and functional analysis of ClbQ, an unusual intermediate-releasing thioesterase from the colibactin biosynthetic pathway. ACS Chem. Biol. 12, 2598–2608 (2017).

Bossuet-Greif, N. et al. Escherichia coli ClbS is a colibactin resistance protein. Mol. Microbiol. 99, 897–908 (2016).

Tripathi, P. et al. ClbS is a cyclopropane hydrolase that confers colibactin resistance. J. Am. Chem. Soc. 139, 17719–17722 (2017).

Colis, L. C. et al. The cytotoxicity of (–)-lomaiviticin A arises from induction of double-strand breaks in DNA. Nat. Chem. 6, 504–510 (2014).

Melvin, M. S. et al. Double-strand DNA cleavage by copper·prodigiosin. J. Am. Chem. Soc. 122, 6333–6334 (2000).

Povirk, L. F., Wübker, W., Köhnlein, W. & Hutchinson, F. DNA double-strand breaks and alkali-labile bonds produced by bleomycin Nucleic Acids Res. 4, 3573–3580 (1977).

Stubbe, J. A. & Kozarich, J. W. Mechanisms of bleomycin-induced DNA degradation. Chem. Rev. 87, 1107–1136 (1987).

Chen, J. & Stubbe, J. Bleomycins: towards better therapeutics. Nat. Rev. Cancer 5, 102–112 (2005).

Piti , M. & Pratviel, G. Activation of DNA carbon–hydrogen bonds by metal complexes. Chem. Rev. 110, 1018–1059 (2010).

Chaturvedi, K. S., Hung, C. S., Crowley, J. R., Stapleton, A. E. & Henderson, J. P. The siderophore yersiniabactin binds copper to protect pathogens during infection. Nat. Chem. Biol. 8, 731–736 (2012).

Humphreys, K. J., Johnson, A. E., Karlin, K. D. & Rokita, S. E. Oxidative strand scission of nucleic acids by a multinuclear copper(II) complex. J. Biol. Inorg. Chem. 7, 835–842 (2002).

Rogakou, E. P., Pilch, D. R., Orr, A. H. Ivanova, V. S. & Bonner, W. M. DNA double-stranded breaks induce histone H2AX phosphorylation on serine 139. J. Biol. Chem. 273, 5858–5868 (1998).

Schultz, L. B., Chehab, N. H., Malikzay, A. & Halazonetis, T. D. p53 binding protein 1 (53bp1) is an early participant in the cellular response to DNA double-strand breaks. J. Cell Biol. 151, 1381–1390 (2000).

Collins, A. The comet assay for DNA damage and repair. Mol. Biotechnol. 26, 249–261 (2004).

Burger, R. M., Peisach, J. & Horwitz, S. B. Activated bleomycin: a transient complex of drug, iron, and oxygen that degrades DNA. J. Biol. Chem. 256, 11636–11644 (1981).

Chan, Y. A. et al. Hydroxymalonyl-acyl carrier protein (ACP) and aminomalonyl-ACP are two additional type I polyketide synthase extender units. Proc. Natl. Acad. Sci. USA 103, 14349–14354 (2006).

Holmes, T. C. et al. Molecular insights into the biosynthesis of guadinomine: a type III secretion system inhibitor. J. Am. Chem. Soc. 134, 17797–17806 (2012).

Stern, B. R. et al. Copper and human health: biochemistry, genetics, and strategies for modeling dose-response relationships. J. Toxicol. Env. Heal. B 10, 157–222 (2007).

Pommier, Y. Drugging topoisomerases: lessons and challenges. ACS Chem. Biol. 8, 82–95 (2013).

Woo, C. M., Li, Z., Paulson, E. K. & Herzon, S. B. Structural basis for DNA cleavage by the potent antiproliferative agent (–)-lomaiviticin A. Proc. Natl. Acad. Sci. USA 113, 2851–2856 (2016).

